# Coping with multiple enemies: pairwise interactions do not predict evolutionary change in complex multitrophic communities

**DOI:** 10.1101/492132

## Abstract

Predicting the ecological and evolutionary trajectories of populations in multispecies communities is one of the fundamental challenges in ecology. Many of these predictions are made by scaling patterns observed from pairwise interactions. Here, we show that the coupling of ecological and evolutionary outcomes is likely to be weaker in increasingly complex communities due to greater chance of life-history trait correlations. Using model microbial communities comprising a focal bacterial species (*Bacillus subtilis*), a bacterial competitor, protist predator and phage parasite, we found that increasing the number of enemies in a community had an overall negative effect on *B. subtilis* population growth. However, only the competitor imposed direct selection for *B. subtilis* trait evolution in pairwise cultures and this effect was weakened in the presence of other antagonists that had a negative effect on the competitor. In contrast, adaptation to parasites was driven indirectly by correlated selection where competitors had a positive and predators a negative effect. For all measured traits, selection in pairwise communities was a poor predictor of *B. subtilis* evolution in more complex communities. Together, our results suggest that coupling of ecological and evolutionary outcomes is interaction-specific and generally less evident in more complex communities where the increasing number of trait correlations could mask weak ecological signals.

## Introduction

Species do not exist in isolation, but rather are embedded in a complex network, linked to other species through a diverse set of trophic and non-trophic interactions [Thompson 2005]. The ecological and evolutionary dynamics of many focal species have been detailed using both theory and empirical approaches, yet we have remarkably little insight into the effects of community complexity on shaping these dynamics [Lytle 2001; Koskella 2014; Barraclough 2015]. Predicting the outcome of these interactions on the density and traits of a given species within a multi-species community is vital for understanding the long-term effects of environmental and biotic change on the biodiversity and stability of communities [e.g. Yoshida 2003; Johnson 2007; Post 2009; Lawrence 2012; Loreau 2013; Donohue 2016; Mrowicki 2016]. Strong, consistent interspecific interactions also have the potential to impose heavy selection pressures on species [Koskella 2011; Lawrence 2012; Friman 2013], with significant implications for evolutionary trajectories [Yoshida 2003]. This predicts that the selection pressures facing an organism in a complex multi-species system are likely to be contingent upon the composition of that community and, moreover, that the addition of species may dampen or promote evolutionary responses to individual selection pressures [e.g. De Mazancourt 2008; Collins 2014; Betts 2015].

Predation [Sherr 2002; Friman 2013; McClean 2015], parasitism [Vos 2009; Penczykowski 2016] and competition [Gause 1934; Hardin 1960; Foster 2012; Friman 2014] are three major sources of antagonistic interactions for organisms. These interactions act to reduce population densities both directly, by increasing mortality and reducing recruitment, and indirectly, through competition for space and resources. How these different types of interactions separately and importantly collectively affect the population densities of focal species and trait evolution is still relatively unexplored. First, it is possible that increasing the number of antagonistic interactions in the community will have additively negative effects on the population density of the focal species. Alternatively, increasing the number of antagonists could weaken effects on focal species if they show negative effects on each other via, for example, intraguild predation [Friman 2016]. Further, the number and type of interactions could also affect the rate and trajectory of focal species evolution. Selection pressures exerted by multiple antagonistic interactions may, for example, result in trade-offs in terms of evolutionary outcomes [Stearns 1989; Friman 2013, 2016]. Adaptations that may be beneficial in one context may be harmful in another, reducing the fitness benefit and, by extension, the rate of evolutionary change [Stearns 1989; Thompson 1994; Friman 2013; Garland 2014]. For example, predation tends to slow down the rate of host-parasite interactions [Friman 2013] and selection by multiple predator species can change the evolutionary trajectory of prey via trade-offs and correlated selection [Friman 2016]. This is because, even though predators [Pernlather 2005; Estes 2011; O’Connor 2015] and parasites [Kutz 2005; Hudosn 2006] exert strong selection for prey survival, the targets and mechanisms under predator and parasite selection are likely to differ. Similarly, evolution of defense against one antagonist might lead to increased susceptibility to another [Friman 2009].

In addition to predatory and parasitic enemies, competitive interactions are a pervasive force of selection and can be indirect or direct, mediated by shared resources or, in the case of bacteria, through antimicrobials [Wang 2017] or bacteriocins [Ghoul 2015]. Recent evidence from bacterial communities suggests that interference competition and parasitism can act synergistically to suppress bacterial growth due to evolutionary trade-offs, where one selection pressure makes focal species more susceptible to the other [Wang 2017]. Furthermore, evolving resistance to strong antagonistic interactions has often been shown to come at a fitness cost [Tollrian 1995; Sheldon 1996; Van Buskirk 2000; Maclean 2004] that might limit the level of resistance or adaptation to the abiotic environment [Scanlan 2015]. While most of the evidence to date suggests that parasites, predators and competitors impose strong selection that often leads to clear fitness trade-offs, trait correlations may also be positive or neutral [e.g. Wright 1999; Ackerly 2007; Chamberlain 2014]. Adaptations selected in one ecological context might therefore have unexpected consequences in other ecological contexts.

Pairwise interactions such as those among competing species have been shown to predict those in more complex communities within single trophic levels [Foster 2012; Rivett 2016]. However, relatively little is known about how different interactions across trophic levels within a community alter evolutionary trajectories. A recent study conducted with multiple predatory protists and a focal bacterial prey suggest that increasing the number of antagonists in the community can weaken pairwise evolutionary dynamics by increasing antagonism between different predator species [Friman 2016]. Similarly, the presence of both specialist and generalist consumers has been shown to change both coexistence and defence evolution in two competing prey species [Hiltunen 2017]. Here, we explore whether predictions from different types of antagonist pairwise interactions scale in communities with increasing complexity. To this end, we manipulated the composition of multitrophic microbial microcosm communities and examined how community complexity moderates the dynamics of a focal bacterium species – *Bacillus subtilis* – and, critically, its ability to evolve resistance to parasites (bacteriophage SPP1) and predators (*Paramecium caudatum*) and to compete for resources with another bacterium (*Serratia marcescens*). We also examined whether exposure to multiple enemies modified *B. subtilis* growth adaptation in abiotic environments in the absence of other species.

We found that increasing the number of enemies in a community had an overall negative effect on *B. subtilis*. However, only competitors imposed direct selection in antagonist monocultures, while both parasites and predators imposed only indirect correlated selection for resistance evolution in antagonist co-cultures. Crucially, ecological and evolutionary outcomes were coupled only under direct selection by competition, whereas no direct ecological or evolutionary signal was observed regarding *B. subtilis* resistance trait evolution. Together, our results suggest that eco-evolutionary outcomes might be interaction-specific and weakly coupled in more complex communities due to increased chance of coincidental life-history trait correlations.

## Methods

### Experimental design

Our experiment consisted of seven sets of species combinations of *Bacillus subtilis* (NCIB3610) with (1) *SPP1* phage parasite in isolation; (2) *Paramecium caudatum*, a generalist [Johnson 1936; Thurman 2010; Banerjii 2015] bacterial predator in isolation; (3) *Serratia marcescens* (ATCC 29632), a competitor of *B. subtilis*, in isolation; (4) *P. caudatum* and *SPP1* phage; (5) *S. marcescens* and SPP1 phage; (6) *S. marcescens* and *P. caudatum;* and (7) *S. marcescens, P. caudatum* and *SPP1* phage, resulting in a total of 7 experimental treatments, each replicated seven times.

### Community assembly

Culture methods followed closely those outlined in McClean *et al*., Lawler and Morin, and Leary *et al*. Microcosms consisted of loosely capped 200 ml glass bottles with 15 g glass microbeads providing spatial habitat structure. The addition of the glass beads allows for species to interact and behave in a more naturally complex environment. Each microcosm received 100 ml medium consisting of one protist pellet (Carolina Biological Supply, Burlington, NC, USA) per 1 litre spring water and two wheat seeds to provide a slow release nutrient source. All media were sterilised before use. Microcosms were maintained at 22 °C and under a 12:12 h light: dark cycle. Nutrients in the microcosms were replenished on day 7 with a replacement of 7 ml of the microcosm volume with sterile medium and one additional sterile wheat seed. *Paramecium caudatum* were obtained from Blades Biological UK. Bacterial strains and SPP1 phage were taken from frozen laboratory stock cultures.

#### P. caudatum washing protocol

The *P. caudatum* protist cultures used for this experiment were laboratory cultures and therefore, while not inoculated with any bacterial populations, were not entirely sterile. To account for this, *P. caudatum* cultures were washed with sterile media before addition to the microcosms to minimise contamination. In addition, the remaining media from the washing process (minus *P. caudatum*) was combined, mixed thoroughly and a similar volume to the protist treatments was added to all microcosms to ensure that any bacteria present in the medium had an equal chance of colonising each microcosm in every treatment. We identified a single bacterial contaminant from this protist media, *Klebsiella sp*., which colonised all microcosm units in this manner in low frequencies and was treated as a standard ‘background’ member of the community [McClean 2015]. At the end of the experiment, the density of the *Klebsiella sp*. did not vary with experimental treatment over the course of the experiment (repeated measures ANOVA; F_2,46_ = 1.23, P = 0.3).

#### Bacterial cultures

Overnight cultures of strains *B. subtilis* (NCIB3610) and *S. marcescens* (*ATCC* 29632), grown in a tryptone yeast (TY) medium (Luria-Bertani broth supplemented with 10 mM MgSO_4_ and 100 M MnSO_4_ after autoclaving [Wach 1996]), were diluted into fresh TY medium at an optical density at 600 nm (OD_600_) ~ 0.03 and grown at 37°C until late exponential phase (OD_600_ ~ 1.0), at which time 1 ml of each bacterial culture was inoculated into 100 ml microcosm medium, as required for experimental treatments. A sample of both the *B. subtilis* and *S. marcescens* cultures were also frozen at -80°C as the ancestral populations for later evolutionary comparisons. Microcosms were left for 24 hr at 37 °C to facilitate growth of the bacteria prior to addition of the phage. 1 ml of a 10^−3^ dilution (in sterile Phosphate Buffer Saline [PBS]) of the phage stock solution (1.7 × 10^4^ pfu ml^−1^) was added to each microcosm as required. Microcosms were then left for a further 24 hr at 37 °C to facilitate bacterial growth and ensure sufficient numbers before the addition of the bacterivorous *P. caudatum*. Microcosms were inoculated with washed *P. caudatum* (approx. 50-70 individuals), as required for experimental treatments and allowed to settle for two days at 22 °C.

#### Community sampling and population density measurements

The point of addition of the predators is considered as Day 0 of the experiment which then ran for ten days. Samples of bacteria, phage and protists were taken every day for the duration of the experiment after carefully homogenising microcosms by shaking. A 0.1 ml sample was taken to count protist numbers using stereo (Olympus SZX9) and compound (Olympus BX60) microscopes. Bacterial densities were measured through direct colony counts (identified by colony morphology) on plates from appropriately diluted samples. Phage numbers were measured through direct plaque counts on plates from appropriately diluted samples. Prior to plating, 3 ml of standard TY medium was inoculated with a stock of ancestral *B. subtilis* and incubated at 37 °C for a minimum of 4 hr or until an OD_600_ of 0.9-1.0 was achieved; 200 βl of the *B. subtilis* culture was added to 10 ml tubes followed by 200 μl of the microcosm sample, mixed gently by hand and incubated at 37 °C for 15 min. Next, 3 ml of soft agar (0.5%) were added to each tube, swirled, and then poured onto preset 1.5% TY agar plates and incubated overnight at 37 °C. The number of plaques on each plate was then counted.

### Measuring evolutionary changes in the life-history traits of focal species B. subtilis

Eight colonies of *B. subtilis* were isolated from each microcosm via agar plating on the final day of the experiment after microcosms were homogenised and vortexed to strip biofilm and ensure representative sampling. The resistance of *B. subtilis* was then assessed against the ancestral populations of SPP1 phage (*n* = 8 populations per microcosm), ancestral *S. marcescens* competitor (*n* = 3 populations per microcosm) and ancestral predatory *P. caudatum* (*n* = 3 populations per microcosm).

#### Competitive ability

The competitive ability of *B. subtilis* from each treatment against the ancestral competitor *S. marcescens* was assessed in direct competition as a deviation from an initial 50:50 abundance ratio in co-culture experiments in 96-well plates containing 200 βl of microcosm medium over a 24 hr period at 22 °C. This was accomplished through counting proportions of the two bacteria based on differences in colony morphologies.

#### Predator defence

The strength of predator defence evolution of *B. subtilis* from each treatment was assessed through biofilm formation in 96 well-plates, which is frequently used as a proxy of bacterial defence strategy based on bacterial cell aggregates that cannot be consumed by protists because they are too large or attached to surfaces [Böhme 2009; Chavez-Dozal 2012; Friman 2015]. To this end, evolved and ancestral *Bacillus* bacteria were grown over 24 hr in 96-well plates containing 200 ml of microcosm medium at 22°C with the addition of approximately 10 washed *P. caudatum* cells. At the end of the 24 hr, *P. caudatum* cell number was counted and biofilm assays were done to test predator defence [Böhme 2009; Chavez-Dozal 2012; Friman 2015] as follows. The liquid medium was decanted and all unattached cells were removed through a water rinse. Next, 100 βl of a 0.1% crystal violet solution was added to the wells to stain the biofilm attached on microplate well walls. The wells were then left for 15 min and rinsed with deionised water. Plates were left to dry overnight before 125 μl of 30% acetic acid (in water) were added to each of the wells and incubated at room temperature for 15 minutes to solubilize the crystal violet. The supernatant from each well was then transferred to wells in a new plate and biofilm production was quantified by measuring the absorbance at 540 nm using 30% acetic acid as the blank.

#### Parasite defence

We used short-term growth assays in liquid media to assess the resistance of *B. subtilis* populations to the ancestral SPP1 phage [Moulton-Brown 2018]. To this end, 10 μl of evolved *B. subtilis* isolates from each treatment was added to 200 μl microcosm media in 96-well plates and allowed to grow independently for 20 hr at 22°C in both the presence and absence of 10 μl of the ancestral SPP1 phage (10^−4^ titre). Bacterial density was then quantified by measuring the OD_600_ to attain a proxy of parasite resistance (the higher the OD, the higher the phage resistance). Resistance was thus measured as reduction in density due to the parasite compared to when the *B.subtilis* populations were grown alone without the parasite.

#### Growth assays of B. subtilis

We compared the growth of evolved and ancestral *B. subtilis* isolates to assess whether the populations from different ecological contexts differed in their adaptation to the growth medium [Scanlan 2015], or if adaptations against different enemies might have incurred costs in terms of trade-offs with growth [Friman 2014; 2015; 2016]. We measured differences in two bacterial growth parameters that are indicative of their ability to compete for resources: maximum growth (population density after 15 hr of growth, by which time they had reached stationary phase), and maximum growth rate (i.e. maximum rate of population growth per hour). Briefly, ancestral and evolved *B. subtilis* isolates were grown independently over a 15-hour period (OD_600_ measured at 10 minute intervals) in the microcosm medium using 96-well plates to assess differences in growth parameters.

### Statistical analyses

All densities were log (x+1) transformed prior to analysis to reduce bias caused by different population sizes. Repeated measures analysis of variance (ANOVA) was used to assess population density changes in each of *B. subtilis, S. marcescens, P. caudatum* and SPP1 phage according to community complexity with post-hoc Tukey honest significant difference (HSD) tests to highlight specific differences among treatments. ANOVA and t-tests were used to examine the evolution of resistance in *B. subtilis* isolates from each of our microcosm communities against each of the competitor (based on density ratios), predator (biofilm density) and parasite (based on density). Finally we used one-sample t-tests to test whether the ecological or evolutionary outcomes from two-species communities could reliably predict the outcomes in more complex communities. Predicted values were calculated as the mean (± s.e.) across antagonist monoculture treatments; For example, the predicted value for the competitor + parasite treatment was estimated by merging the competitor alone and parasite alone treatments and calculating their combined mean. All analyses were carried out in R (version 3.4.4; R Developmental Core Team 2018). All data available as supplementary files.

## Results

### Population density dynamics of B. subtilis focal species

The density of *B. subtilis* varied significantly with the number of enemy species present in the community (repeated measures ANOVA: *F*_2,46_ = 8.07, *P* < 0.01 ; Figure 1). Though addition of a second enemy did not alter the density of *B. subtilis* (Tukey contrasts: 2 enemies present – 1 enemy present; *P =* 0.25; Figure 1), regardless of antagonist identity (Table S1), *B. subtilis* densities were significantly lower when all three antagonistic species were present together in the community (Tukey contrasts: 3 enemies present – 1 enemy present; *P* < 0.05, 3 enemies present – 2 enemies present; *P* < 0.05; Figure 1). The observed density of *B. subtilis* in multispecies communities was consistently lower than that predicted based on densities observed in two species co-cultures (Table S1; Figure 3a). Together, these results demonstrate that pairwise communities did not predict the density regulation effects observed in the three-enemy community.

**Figure 1.**
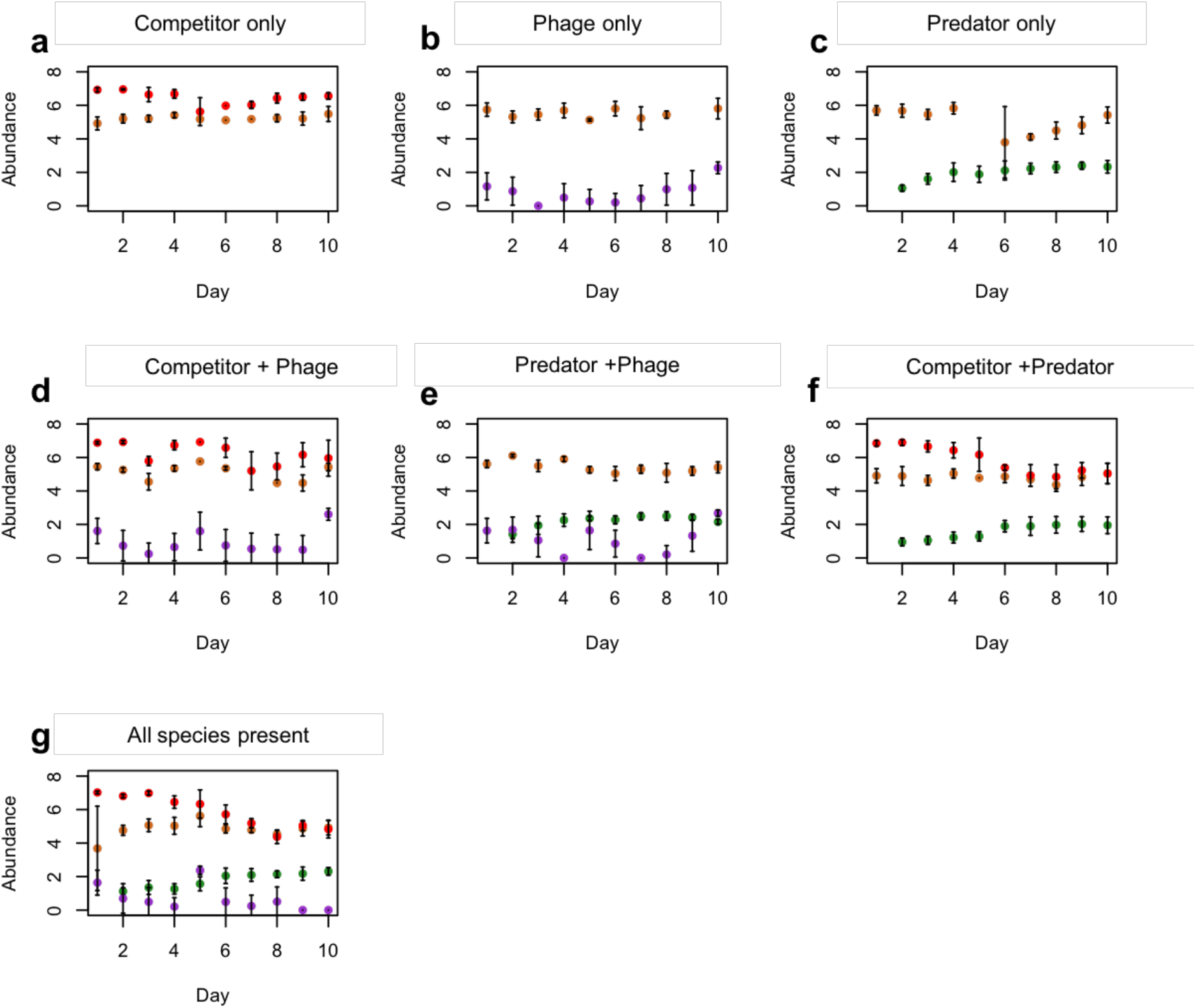
Abundance (log-transformed, mean ± std. dev., *n* = 7) of the focal species *B. subtilis* (orange circles), competitor bacterium *S. marcescens* (red circles), predator *P. caudatum* (green circles) and *B. subtilis-specific* parasite phage SPP1 (purple circles) in each treatment over the duration of the experiment (Days 1-10).

### Population density dynamics of competitor, parasite and predator

The bacterial competitor species *S. marcescens* reached higher population densities compared to *B. subtilis* when co-cultured in the absence of other antagonists (Figure 1a). This is indicative of competitive advantage. However, when either the specialist parasite of *B. subtilis* (phage SPP1), a generalist predator species (*P. caudatum*), or both, were added to the microbial community, this competitive advantage was diminished, as evinced in a greater reduction in *S. marcescens* relative to *B. subtilis* densities (repeated measures ANOVA: F_2, 25_ = 10.34, *P* < 0.001; Tukey contrasts; 2 enemies – 1 enemy and 3 enemies – 1 enemy, *P* < 0.05 in all cases; Figure 1). This suggests that the presence of parasites and predators, either separately or together, evened out the competitive difference between the two competing bacterial species. Parasite densities varied with community richness (repeated measures ANOVA: *F*_2, 25_ = 4.1, *P* < 0.05). Parasite density was unaffected by the presence of other antagonists (Tukey contrasts: 3 enemies – parasite, *P* = 0.8; 2 enemies–parasite, *P* = 0.1; Figure 1b, g), except in the presence of both the competitor and predator of *B. subtilis*, which decreased parasite density relative to either species present in isolation (Tukey contrasts: 3 enemies – 2 enemies, *P* < 0.05; Figure 1). Parasite densities dropped below the detectable limit by the end of the experiment (Figure 1g) in the presence of both competitor and predator (Tukey contrasts: All species present – every other parasite treatment, *P* < 0.05 in all cases; Figure 1). This suggests that the competitor and predator had relatively small effects on parasite densities in communities containing two enemies but additive negative effects in three-antagonist communities.

Although predator densities in general increased over time, they did not vary with antagonist richness (repeated measures ANOVA, *F*_2,25_ = 0.63, *P* = 0.5; Figure 1). Thus, the predator was the least affected by the presence of other interacting antagonistic species in the community.

### Evolution of B. subtilis competitive ability and resistance against parasite and predator

Evolutionary change in the competitive ability of *B. subtilis* was assessed by comparing the growth of ancestral and evolved *B. subtilis* clones in direct competition with the ancestral competitor species, *S. marcescens*, at the end of the experiment. We found that the competitive ability of *B. subtilis* increased only if it had been exposed to *S. marcescens* in the absence of other antagonists (Figure 2a; ANOVA: *F*_6, 19_ = 4.58, *P* = 0.005; Tukey contrasts; *P* < 0.05 for each treatment compared to *B. subtilis* + *S. marcescens* alone, except for *B. subtilis* + SPP1 phage parasite, where *P* = 0.08). The presence of any other enemy limited the evolution of *B. subtilis* competitive ability and no difference was observed relative to the ancestral strain (Tukey HSD; *P* > 0.05 between all other treatment pairs, Figure 2a). The observed changes in *B. subtilis* competitive ability were lower than predicted based on pairwise co-cultures, except for the predator-parasite treatment (Table S1; Figure 3b). These results therefore indicate that evolutionary changes in *B. subtilis* competitive ability were weakened in the presence of additional species.

**Figure 2.**
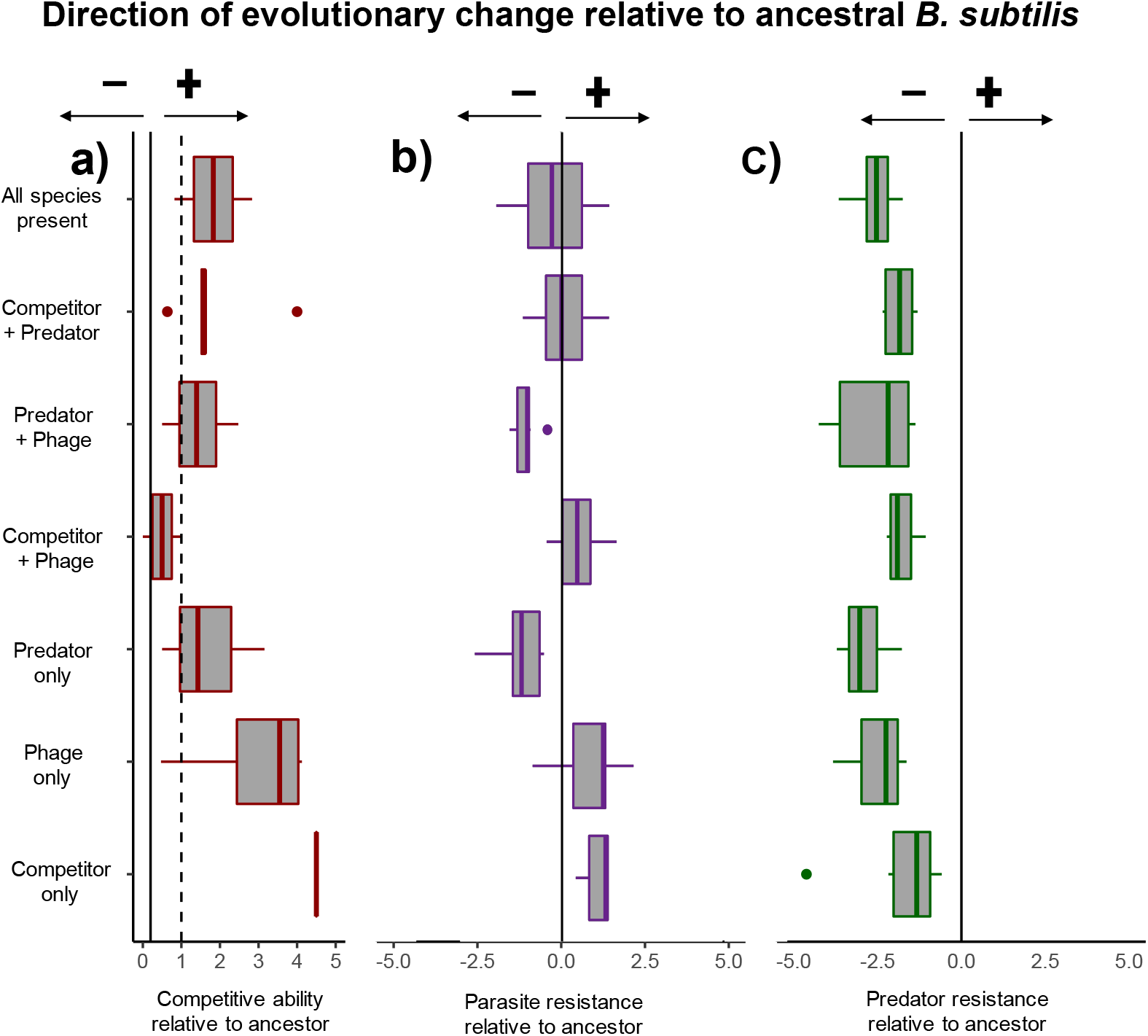
The effect of community composition on *B. subtilis* evolution relative to the ancestral strain (solid line). **a**. Competitive ability of *B. subtilis* isolates, measured as the ratio of *B. subtilis* to *S. marcescens* (dashed line indicates a 1:1 ratio, solid line represents ancestral resistance). **b**. Relative resistance to parasite in *B. subtilis* across treatment groups. Resistance to phage SPP1 was measured as the difference in growth (optical density) of *B. subtilis* in the presence or absence of the ancestral phage parasite. The solid line represents the resistance of ancestral *B. subtilis* and the resistance of our experimental treatment isolates relative to ancestral parasite resistance. **c**. Growth of *B. subtilis* isolates in the presence of the predator *P. caudatum* (measured as log_10_ optical density of biofilm production). Solid line indicates resistance of the ancestral *B. subtilis* grown in the presence of *P. caudatum*). In all cases, values to the right of the solid lines indicate higher relative resistance than the ancestral strain, values to the left of the solid line indicate lower resistance relative to ancestral strain of *B. subtilis*.

**Figure 3.**
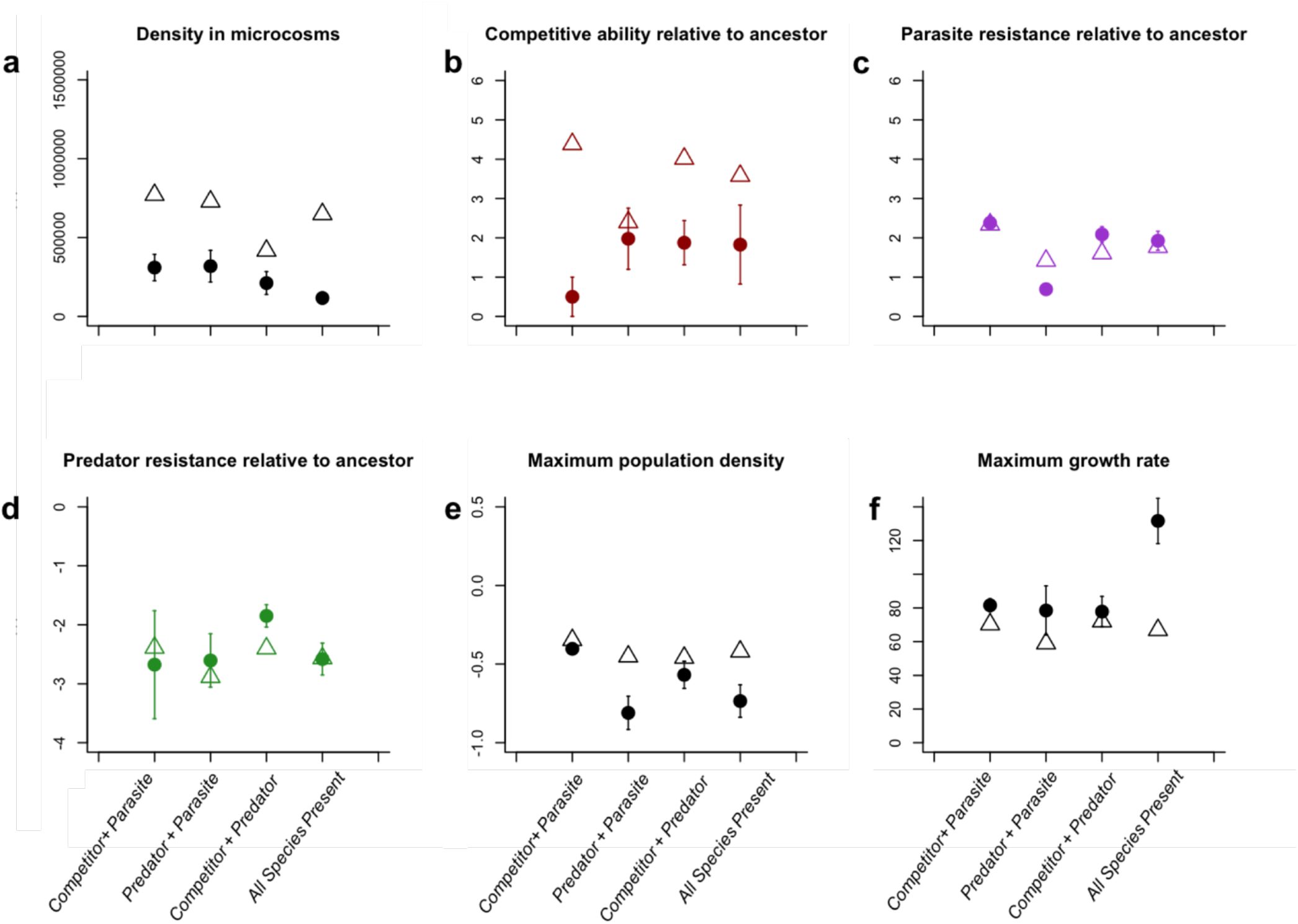
Predicted (open triangles) and observed (closed circles) measurements of *B. subtilis* in multispecies communities**. a**. Density of *B. subtilis* per 1 ml microcosm medium. **b**. Maximum population density. **c**. Maximum growth rate of evolved isolates **d**. Competitive ability of *B. subtilis* isolates, measured as the ratio of *B. subtilis* to *S. marcescens*. **e**. Relative resistance to parasites of *B. subtilis* across treatment groups. **f**. Growth of *B. subtilis* isolates in the presence of the predator *P. caudatum*. Predicted values were calculated as the mean (± s.e.) across antagonist monoculture treatments; For example, the predicted value for the competitor + parasite treatment was estimated by merging the competitor alone and parasite alone treatments and calculating their combined mean.

Evolution of resistance to the parasite was measured as the difference in the growth of ancestral and evolved *B. subtilis* isolates in the presence and absence of the ancestral SPP1 phage (Figures 2b, S1). Overall, resistance to the ancestral parasite varied strongly depending upon community composition (ANOVA: *F*_6, 38_ = 7.46, *P* < 0.001; Figure 2b). Prior exposure to the parasite alone did not affect *B. subtilis* resistance to the ancestral parasite (*t*-test; *t*_8.14_ = 0.98, *P* = 0.35; Figure 2b, S1). In contrast, the correlated evolutionary response to *S. marcescens* alone led to high ancestral parasite resistance (*t*-test; *t*_7.3_ = 5.1, *P* = 0.001; Figures 2b, S1), while this effect disappeared with the addition of other antagonists (*t*-tests; parasite: *t*_11.03_ = 0.57, *P* = 0.58; predator: *t*_9.9_ = 0.74, *P* = 0.48; Figures 2b, S1). Moreover, prior exposure to the predator alone (*t*-test; *t* _8.89_ = -3.7, *P* < 0.01 ; Figures 2b, S1), or in combination with the phage parasite (*t*-test; *t*_9.52_ = 8.58, *P* < 0.001 ; Figures 2b, S1), reduced *B. subtilis* resistance to the parasite. When all three enemy species were present together, there was no effect on *B. subtilis* resistance to the ancestral SPP1 parasite (*t*-test; *t*_9.99_ = 1.27, *P* = 0.23; Figures 2b, S1). No major differences were observed between predicted and observed parasite resistance values largely because only weak parasite resistance was observed in general (Figure 3). However, when both predator and parasite were present together, the resistance of *B. subtilis* was lower than expected based on pairwise co-culture predictions (Table S1; Figure 3c). Together, these results suggest that both competitors and predators exerted opposing, correlated selection for *B. subtilis* parasite resistance and that they cancelled the effect of each other out in competitor-predator co-cultures.

Resistance to the predatory protist *P. caudatum* was measured as the difference in the extent of biofilm formation between ancestral and evolved *B. subtilis* isolates in the presence and absence of a stock culture of *P. caudatum*. As expected, all isolates produced a greater amount of biofilm in the presence of the predator (Figures 2c, S2). However, all evolved isolates produced less biofilm than the ancestral *B. subtilis* strain in the presence of the predator (grey line on Fig 2c), and no difference in biofilm formation was observed between evolved isolates from different antagonist communities (ANOVA: *F*_5, 34_ = 0.77, *P* = 0.58; Figures 2c, S2). As a result, only weak predator resistance evolution and a good agreement between predicted and observed evolutionary outcomes was observed, except for the competitor-predator treatment, where observed evolutionary changes were less than those predicted (Table S1; Figure 3d).

### Determining B. subtilis adaptation to the growth media and the cost of resistance

We quantified the growth of ancestral vs. evolved *B. subtilis* isolates to assess whether populations from different antagonist communities differed in their adaptation to the growth medium and to identify the potential cost of resistance. None of the evolved selection lines showed improved growth relative to the ancestral *B. subtilis* strain, which suggests that no adaptation to the growth media occurred and that changes are likely due to some costs of resistance (Figure S3a). The maximum population density of evolved *B. subtilis* isolates was unaffected by previous evolutionary history with either the competitor *S. marcescens* or parasite (ANOVA; main effect of *S. marcescens: F*_1,38_ = 0.64, *P* = 0.43; main effect of SPP1 phage: *F*_1,38_ = 1.5, *P* = 0.23; Figure S3a). The maximum population density of *B. subtilis* was, however, reduced significantly if it had previously been exposed to antagonist communities that included the predator (ANOVA; main effect of *P. caudatum: F*_1,38_ = 36.12, *P* < 0.001; Figure S3a), and this reduction was the strongest in the predator-parasite treatment. In line with this, the observed maximum densities were lower than predicted in more complex communites that included the predator (Table S1; Figure 3e).

The growth rate of evolved *B. subtilis* isolates was much lower compared to the ancestral strain (Figure S3b) and only *B. subtilis* that had evolved in the presence of all three enemies had growth rates comparable with the ancestral strain. In general, reduction in *B. subtilis* growth rate was lower in antagonist communities that included the *S. marcescens* competitor species (ANOVA; main effect of *S. marcescens: F*_1,38_ = 16.44, *P* < 0.001; Tukey contrasts *P* < 0.05; Figure S3b) and *P. caudatum* predator (ANOVA; main effect of *P. caudatum: F*_1,38_ = 8.21, *P* = 0.007; Tukey contrasts *P* < 0.05; Figure S3b). In contrast, the presence of the SPP1 phage parasite had no clear effect on *B. subtilis* growth rate (ANOVA; main effect of SPP1 phage: *F*_1,38_ = 2.54, *P* = 0.12; Figure S3b). Due to similar evolutionary trajectories in pairwise co-cultures, good agreement was found between predicted and observed evolutionary outcomes (Figure 3). However, the observed reduction in *B. subtilis* growth rate was much less than predicted in the three-antagonist community (Table S1; Figure 3f). Together, these results suggest that evolved isolates had generally lowered maximum growth and growth rates compared to the ancestral strain and that these changes were driven mainly by the presence of the predator.

## Discussion

In this study, we set out to test whether pairwise interactions observed both within and among trophic levels scale with increasing complexity in multitrophic microbial communities. We found that, while increasing the number of enemies in a community had an overall negative effect on the densities of our focal bacterium, *B. subtilis*, only the competitor imposed direct selection for *B. subtilis* trait evolution in pairwise cultures. Further, this effect was weakened in the presence of other antagonists that had a negative effect on the competitor. As a result, ecological (population density) and evolutionary (trait evolution) outcomes were coupled only in the case of *B. subtilis* competitive ability. However, both predators and parasites affected *B. subtilis* trait evolution indirectly in antagonist co-cultures, having either positive or negative effects respective to the ancestral strain. In case of both ecological and evolutionary outcomes, selection in pairwise communities was a poor predictor of outcomes in more complex communities, and often led to overpredictions. Overall, our findings indicate that the coupling of eco-evolutionary outcomes is both trait- and interaction-specific and harder to predict in more complex communities where the increasing number of trait correlations can mask weak ecological signals and alter predictions based on pairwise co-cultures.

Only the competitor bacterium *S. marcescens* selected for clear evolutionary change in *B. subtilis* in single-enemy monocultures. Under experimental conditions, *S. marcescens* attained higher densities than *B. subtilis*, indicating a competitive advantage. This manifested as a strong driving force for evolutionary change in *B. subtilis*, where isolates from the competitor monoculture treatment demonstrated much greater competitive ability towards their ancestral competitor in comparison to both ancestral *B. subtilis* and evolved *B. subtilis* isolated from multi-enemy co-cultures. In contrast, neither the parasitic phage nor predatory protist caused such a magnitude of evolutionary change against ancestral enemies in *B. subtilis*. One explanation for this is that the duration of the experiment (10 days) was not sufficiently long for the resistance to sweep through *B. subtilis* bacterial populations and become fixed. Also, the nutrient concentration of our experimental conditions was relatively low, which has been shown previously to slow down both parasite [Friman 2015] and predator [Friman 2011] resistance evolution. In the case of predator resistance evolution, we observed a very strong plastic defence response, which decreased with evolved compared to ancestral *B. subtilis* strains. While this is indicative of defence evolution, it was uniform among antagonist treatments, suggesting that it was perhaps driven independent of the presence of the predator (*e.g*. adaptation to growth media). It is less clear why no clear signs of parasite resistance were observed. It is possible that *B. subtilis* evolved resistance to its contemporary parasite without clear effects on resistance to the ancestral parasite. Also, phage densities were quite low in our experimental system, which could also have reduced the likelihood of resistance evolution [Lopez-Pascua 2008; Friman 2008].

The observed competitive advantage and selective force of the competitor bacterium on *B. subtilis* was weakened substantially in the presence of other enemies. The presence of either predator or parasite, alone or in combination, led to a reduction in *S. marcescens* density, negating the effect of the competitor on driving selection in *B. subtilis* competitive ability. Alternatively, it is possible that selection by parasites or predators somehow conflicted with the selection by competitors. For example, the presence of multiple enemies can weaken the respective effects via trade-offs where selection by one antagonist makes the focal species more susceptible to the other antagonists [Friman 2009]. This is, however, an unlikely explanation in our study, as selection by the competitor alone did not make *B. subtilis* more susceptible to the predator or parasite.

We found no evidence of evolutionary change in *B. subtilis* response to either predator or parasite in monocultures. In contrast, adaptation to the parasitic bacteriophage was driven indirectly by selection in multispecies communities, where the presence of competitors had a positive, and predators a negative, effect. The density of parasites was lowest in the presence of both other antagonists but this does not appear to have altered the strength of their evolutionary selective pressure on *B. subtilis*. This implies that adaptations conferring a competitive advantage against *S. marcescens* also conferred a level of resistance against bacteriophage SPP1. There are several potential mechanisms through which bacteria may develop resistance against both a competitor and a bacteriophage parasite simultaneously. For instance, the production of extra-cellular compounds – including matrices and competitive inhibitors— are known to confer both competitive advantages [Hibbing 2010] and resistance [Labrie 2010] against host-specific bacteriophage parasites. Only plastic defensive response to predation through biofilm production was observed and this was not affected by antagonist community composition. When both competitor and predator were present together, with or without the parasite, the resistance of *B. subtilis* to the ancestral parasite was similar to that of the ancestral *B. subtilis*. As the predator appeared to feed non-preferencially on all bacteria present, it had a negative effect on the densities of both *B. subtilis* and *S. marcescens*, which likely evened out some of the competitive interactions between the two species. In addition, it is likely, though not directly measured here, that predation selected for *S. marcescens* defence evolution which may have altered competitive interactions between the bacteria [Hiltunen 2017]. Together, these results suggest that competitors and predators impose opposing selection but that, when present in the community together, cancel each other out indicative of cryptic evolution [Yoshida 2007].

Our focal bacterium, *B. subtilis*, adapted differentially to the growth media depending on the species that it was co-cultured with. The presence of single enemies did not increase or reduce the maximum growth obtained by the *B. subtillis* populations relative to their ancestral strain. This implies that changes in maximum growth would likely be due to costs of resistance against multiple enemies. However, the presence of predator and phage together, with or without the competitor, led to lowered maximum growth. This suggests that the predators and parasites had additive negative effects on the growth of the bacterium. We also found that all populations of single-or two-enemy communities had lower growth rates than the ancestral strain. In contrast, the presence of all three enemies at once led to increased growth rates in the growth medium relative to the other populations. It has been shown previously that evolution of defencive traits can impose costs in terms of reduced growth rates [Ford 2016; Ashby 2017]. The effects of community complexity on a given species depend upon the growth environment that they find themselves in, how each species is adapted to it and, thus, the relative costs of defencive resistance evolution. However, we found that resistance evolution did not increase clearly with community complexity and is thus unlikely to explain the lowest growth rate cost in the most complex community.

Neither the ecological nor the evolutionary outcomes of our focal bacterium *B. subtilis* could be predicted reliably based on pairwise interactions. We found that observed densities of *B. subtilis* populations in complex communities were consistently lower than predicted from pairwise cultures across all treatments. This is suggestive of synergistic interactions among enemy communities. Similarly, the evolutionary outcomes of *B. subtilis* were more varied and often in disagreement between observed and predicted outcomes in general. However, these mismatches depended upon the composition of specific enemy communities and the given traits measured. Most of the discrepancies between predicted and observed evolutionary outcomes were associated with competitive ability and the growth parameters of our evolved isolates, while no clear pattern was found with increasing community complexity. The greatest magnitude of evolutionary response in monoculture was observed with the compeitior bacterium S. marcescens, an effect which was no longer observed with increasing complexity. This could suggest that the greater the magnitude of pairwise interactions, the greater the extent that increased complexity will modify that response in natural communities.

The coupling of eco-evolutionary outcomes were also trait-specific: evolutionary changes in the competitive ability of *B. subtilis* were linked strongly with ecological dynamics, whereas the evolution of parasite resistance did not exert a clear ecological signal. When species are dealing with a single or few strong interactions it might be expected that ecological dynamics are tightly coupled with evolutionary trajectories, such that relative species densities may indicate the strength of selection. Ecological parameters such as biomass are frequently used to elucidate shifts in evolutionary outcomes (such as changes in the ability to resist parasites) within populations [Yoshida 2007]. Shifts in a population’s evolutionary strategy in terms of growth rates and resistance to enemies have the potential to greatly alter the ecological interactions within a community. Detecting such evolutionary signals of change within a population in different ecological contexts is of great importance for predicting the impact of changing environments and biodiversity loss. When considering the impact of global change on species evolution, our results suggest that competitive interactions may play a far stronger role than appreciated previously. Competition in microbial communities is a known driver of many species traits [Schutler 2015; Ashby 2017; Niehus 2017] and our study shows that exploitative competitive interactions may be even more important for predicting responses to other biotic stressors than strong trophic interactions.

In conclusion, our findings demonstrate that predicting ecological and evolutionary dynamics of complex systems based on pairwise interactions is extremely challenging. If we are to better predict eco-evolutionary outcomes in large multispecies ecosystems, we must endeavor to scale up empirical experiments to incorporate a greater range of more realistic interaction networks, incorporating multiple trophic levels. Moreover, our results suggest that selection in one context can have unexpected consequences for species interactions in another context due to negative and/or positive trait correlations. It is thus crucial to try to understand targets of selection more mechanistically in order to predict how different selection pressures might shape species evolution in complex communities. Incorporating these approaches into current eco-evolutionary dynamics frameworks will help to achieve much greater insight into the evolution and mechanisms of trait correlations and, therefore, enhance our capacity to understand and predict species response to global change in complex communities.

## Supporting information

