## Supplementary figures and images for "Coping with multiple enemies: pairwise interactions do not predict evolutionary change in complex multitrophic communities"

### Supplementary file 1

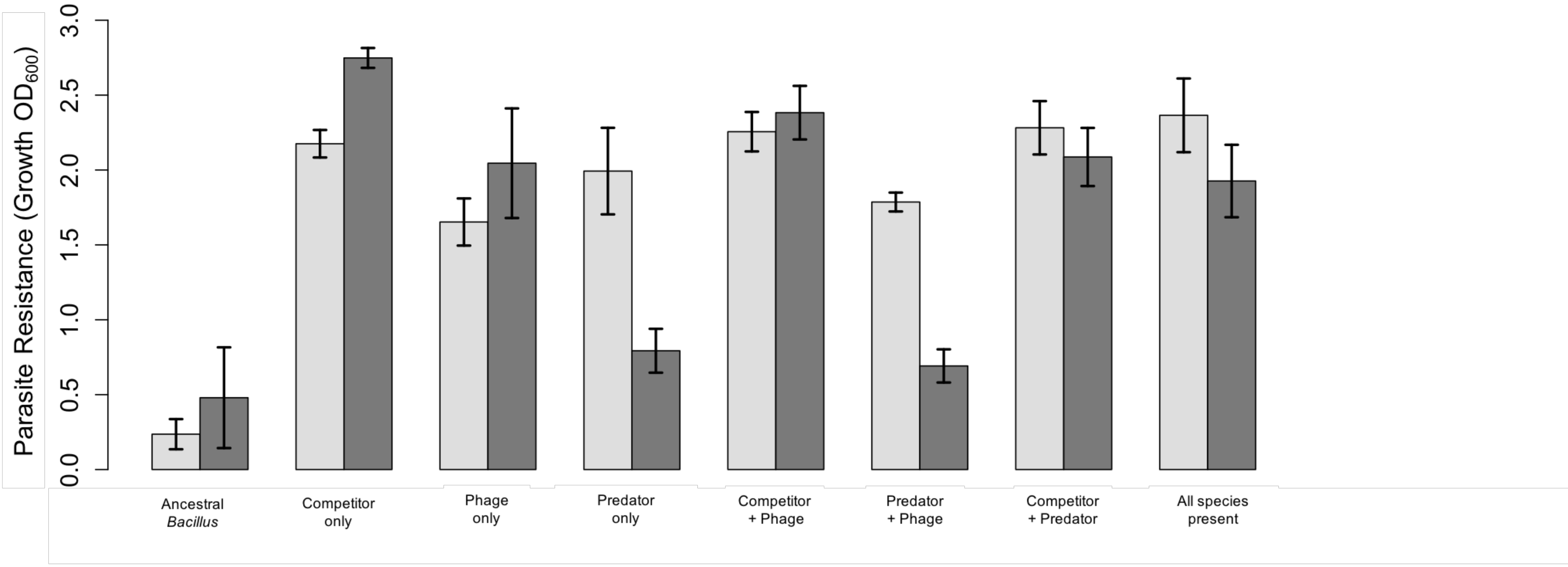

### Supplementary file 2

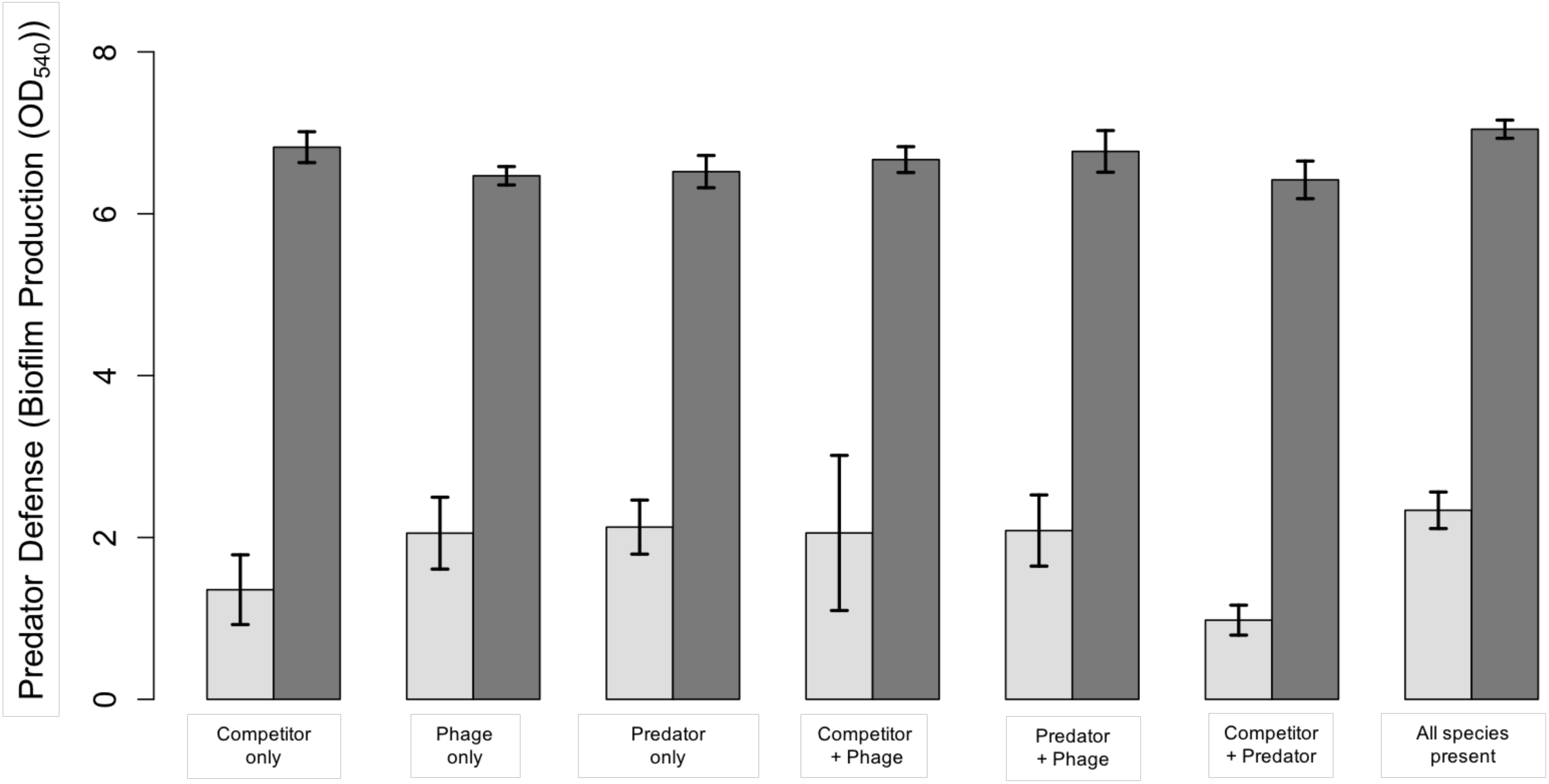

### Supplementary file 3

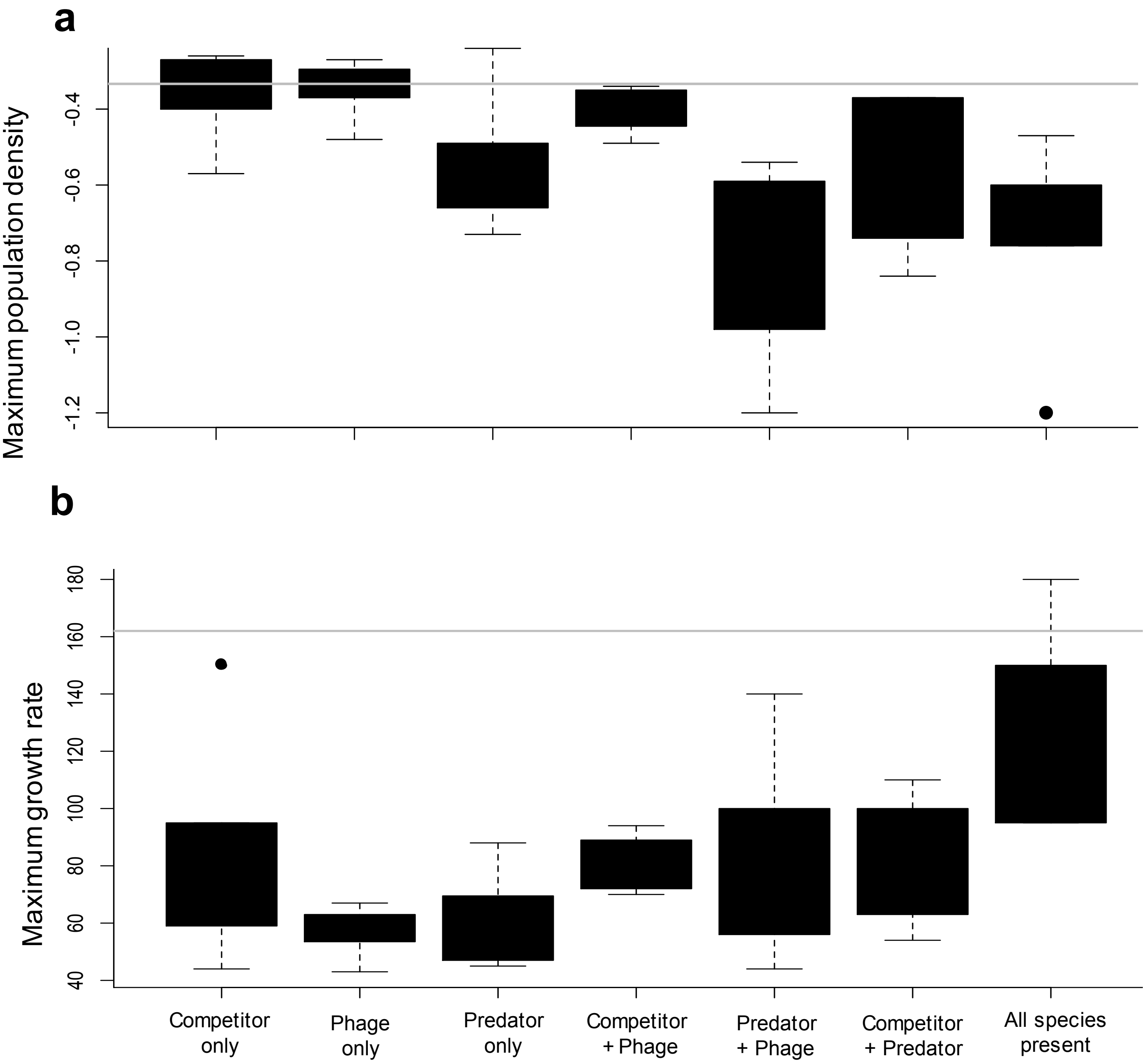
