## Supplementary material for "Coping with multiple enemies: pairwise interactions do not predict evolutionary change in complex multitrophic communities"

| | Treatment | Df | t | P |
|---|---|---|---|---|
| ***Density in Microcosms*** | Competitor + Parasite | 6 | -5.48 | **0.002** |
| | Competitor +Predator | 6 | -2.84 | **0.03** |
| | Predator + Parasite | 4 | -4.07 | **0.02** |
| | Full Community | 5 | -14.25 | **< 0.001** |
| ***Competitive Ability*** | Competitor + Parasite | 4 | -17.58 | **< 0.001** |
| | Competitor +Predator | 4 | -3.81 | **0.02** |
| | Predator + Parasite | 6 | -0.7 | 0.51 |
| | Full Community | 5 | -5.88 | **0.002** |
| ***Parasite Resistance*** | Competitor + Parasite | 6 | 0.25 | 0.81 |
| | Competitor +Predator | 5 | 2.47 | **0.056** |
| | Predator + Parasite | 6 | -6.56 | |
| | Full Community | 5 | 0.65 | 0.54 |
| ***Predator Defense*** | Competitor + Parasite | 6 | -0.32 | 0.76 |
| | Competitor +Predator | 5 | 2.9 | **0.03** |
| | Predator + Parasite | 6 | 0.62 | 0.55 |
| | Full Community | 5 | -0.05 | 0.96 |
| ***Maximum Population Density*** | Competitor + Parasite | 6 | -2.49 | 0.05 |
| | Competitor +Predator | 5 | -1.26 | 0.26 |
| | Predator + Parasite | 5 | -3.39 | **0.02** |
| | Full Community | 5 | -3.05 | **0.03** |
| ***Maximum Growth Rate*** | Competitor + Parasite | 6 | 2.92 | **0.03** |
| | Competitor +Predator | 5 | 0.65 | 0.55 |
| | Predator + Parasite | 5 | 2.088 | 0.09 |
| | Full Community | 5 | 4.81 | **0.005** |
